# Distinct *cis*-acting elements govern purine-responsive regulation of the *Leishmania donovani* nucleoside transporters, LdNT1 and LdNT2

**DOI:** 10.1101/2020.10.15.336420

**Authors:** M. Haley Licon, Francesca Goodstein, Diana Ortiz, Scott M. Landfear, Phillip A. Yates

## Abstract

Purine salvage from the host is an obligatory process for all protozoan parasites. In *Leishmania donovani*, this is accomplished by four membrane nucleoside and nucleobase transporters, or LdNTs. Previously, we demonstrated that purine starvation invokes a robust stress response in *Leishmania* and characterized the proteomic changes involved. However, because *Leishmania* do not control the transcription of individual genes, the underlying mechanisms responsible for these changes were ill-defined. LdNT1 and LdNT2 are among the most rapidly and significantly upregulated genes in purine-starved *L. donovani* parasites. Thus, to better understand post-transcriptional mechanisms of purine-responsive gene expression, we have examined regulation of these genes in molecular detail. Here we report that LdNT1 and LdNT2 are controlled by distinct *cis*-acting elements. In the case of LdNT2, mRNA abundance and translational enhancement under purine stress depend on a 76 nt-long polypyrimidine tract encoded in the *LdNT2* mRNA 3’-UTR. Transcripts containing the *LdNT2* polypyrimidine tract were found to localize to discrete cytoplasmic foci in purine-replete cells, suggesting that the *LdNT2* message may be stored in RNA granules at steady-state. In the case of LdNT1, we found that purine-responsiveness is conferred by a 48 nt-long polypyrimidine tract and additional upstream element, termed UE1. Both features are independently required for regulation, with the polypyrimidine tract and UE1 controlling mRNA abundance and translation, respectively. Together, these results highlight a remarkable degree of complexity in the regulation of the *Leishmania* purine stress response and set the stage for future investigations to identify the larger network of RNA-protein and protein-protein interactions involved.

## Introduction

Kinetoplastid parasites from the *Leishmania* genus are the causative agents of leishmaniasis, a suite of debilitating and life-threatening diseases that disproportionately affect the developing world [WHO, 2010]. Throughout their lifecycles, *Leishmania* transition between flagellated promastigote forms in the midgut of a sandfly vector and intracellular amastigotes, which survive and replicate within the macrophages of a vertebrate host. These compartments differ significantly in terms of pH, temperature, and nutrient availability, and each presents a unique set of challenges that the parasites must overcome for colonization [Dostálová, 2012; Zilberstein, 1994; Moradin, 2012]. Consequently, mechanisms of stress tolerance and adaptation are integral to progression of the *Leishmania* lifecycle.

Adapting to environmental change requires the activation of specific stress-response gene networks. For prokaryotes and eukaryotes alike, this typically begins in the nucleus with mRNA synthesis. However, *Leishmania* and related kinetoplastids have evolved to bypass transcriptional control entirely. In these organisms, transcription by RNA polymerase II is polycistronic in nature, resulting in the production of multi-gene ‘pre-mRNAs’ [Ivens, 2005]. Individual messages are subsequently produced from this precursor through *trans*-splicing of a conserved, capped exon sequence (the spliced leader) to the 5’ end of each message and 3’ polyadenylation [reviewed in Clayton, 2019]. Consequently, post-transcriptional mechanisms such as mRNA stability and translation have been elevated as key determinants of gene regulation in *Leishmania*. As with higher eukaryotes, these processes are governed primarily by the interactions of *cis*-acting elements in mRNA and *trans*-acting RNA-binding proteins (RBPs) [Kolev, 2014]. Considerable effort has therefore been devoted to the study of such elements in kinetoplastids; however, few have been implicated in the context of specific stress-response pathways [Droll, 2013; Minia, 2016].

Unlike their insect and vertebrate hosts, *Leishmania* are unable to synthesize purines *de novo* and must scavenge these nutrients from the extracellular milieu [Boitz, 2014]. In *Leishmania donovani*, this is accomplished by the activity of four dedicated nucleoside and nucleobase transporters, or LdNTs, which function as proton symporters to concentrate purines inside the cell [Landfear, 2004]. Despite this dependence, purine stress appears to factor regularly into the *Leishmania* lifecycle and *Leishmania* have evolved a robust stress response to cope with periodic purine scarcity [Carter, 2010; Martin, 2014]. Indeed, purine restriction is required for efficient metacyclogenesis within the sandfly, suggesting that this particular stressor may serve as an important trigger for lifecycle progression [Serafim, 2014]. Purine starvation is readily induced *in vitro* and we previously demonstrated that, upon removal of purines from the culture medium, *L. donovani* promastigotes cease replication and exhibit a characteristic elongation of the cell body [Carter, 2010]. At the same time, the *L. donovani* proteome is dramatically restructured to enhance purine salvage and recycling, minimize energy expenditure, and increase general stress tolerance [Martin, 2014]. Proteins with annotated functions in nucleic acid metabolism and translation are dramatically reduced, consistent with parasites exiting the cell cycle [Martin, 2016]. In addition, a *Leishmania* homolog of the eIF4E cap-binding protein was recently found to concentrate in ribosome-containing cytoplasmic granules under purine stress, consistent with a mechanism of translational repression [Shrivastava, 2019]. However, the molecular mechanisms underlying purine-stress tolerance remained largely unknown.

Three of the four membrane purine transporters, namely LdNTs 1-3, are among the earliest and most significantly upregulated proteins in purine-starved *L. donovani* promastigotes. Thus, to investigate mechanisms of purine-responsive expression, we have used these genes as a model. In previous work, we compared the RNA-centric processes affecting LdNT regulation [Martin, 2014; Soysa, 2014]. LdNTs 1-3 display robust translational enhancement under purine stress, despite a global reduction in protein synthesis [Martin, 2014; Shrivastava, 2019]. At the transcript level, both *LdNT1* and *LdNT3* are increased, whereas *LdNT2* is either unchanged or modestly reduced. Interestingly, though all four of the LdNTs are encoded by relatively high-copy messages, *LdNT2* is exceptional in its steady-state abundance, ranking in the 99.8^th^ percentile in log-stage *L. donovani* promastigotes [Martin, 2014]. Together, these differences suggest that the purine transporters are regulated by independent, though possibly intersecting, post-transcriptional mechanisms.

We recently published that LdNT3 is controlled by a purine-responsive stem-loop in the mRNA 3’-UTR, which operates through mRNA destabilization and translational repression to restrict expression under purine-replete conditions [Licon, 2020]. This element was highly conserved in the orthologous genes from other parasitic kinetoplastids but was not found elsewhere in the *L. donovani* genome. Now we describe additional efforts to characterize the *cis-* acting sequences controlling nucleoside transporters LdNT1 and LdNT2. By systematic deletion mutagenesis, we identified a 76 nt-long polypyrimidine tract in the *LdNT2* mRNA 3’-UTR. Loss of this region led to a drastic reduction in transcript abundance and prevented translational enhancement under purine stress. Additionally, transcripts containing the *LdNT2* polypyrimidine tract localized to discrete cytoplasmic foci in purine-replete cells, suggesting that the high-copy *LdNT2* message may be stored in RNA granules at steady-state. In the case of LdNT1, we found that purine-responsiveness is conferred by a polypyrimidine tract and additional upstream element, termed UE1. We established that both features are independently required for regulation, with the polypyrimidine tract and UE1 controlling mRNA abundance and translation, respectively. Finally, we found that the *LdNT1* polypyrimidine tract can substitute for that of *LdNT2* to confer regulation in the context of the *LdNT2* 3’-UTR.

## Results

### Deletional mutagenesis of the LdNT2 UTRs reveals that purine-responsive expression is mediated by a 76 nt-long polypyrimidine tract

To better understand regulation of the nucleoside transporters in *Leishmania*, we began with a molecular dissection of *LdNT2*. It is published that a luciferase reporter flanked by the *LdNT2* 5’- and 3’-UTRs is endowed with purine-responsive expression, strongly implicating the presence of distinct regulatory sites within one or both of these regions [Martin, 2014]. However, neither UTR was tested independently. As most *cis*-acting RNA elements have been identified in 3’-UTRs [Clayton, 2019], we suspected the *LdNT2* 5’-UTR is dispensable for purine-responsive expression. To test this, we used the approach depicted in Figure 1A. A NanoLuciferase reporter construct (heretofore referred to as *LdNT2/NLuc*) was expressed from the endogenous *LdNT2* locus under the control of either wildtype *LdNT2* UTRs or a 5’-UTR from a purine-unresponsive control gene. For this purpose, we employed nucleobase transporter LdNT4, which, though functionally analogous to LdNTs 1-3, is not differentially regulated with respect to purines [Martin, 2014]. The 5’-UTRs were substituted so as to preserve the dominant *LdNT2* 5’ splice-acceptor site, located 269 nt upstream of translation start [TriTrypDB.org]. As we previously established that the *LdNT2* CDS is itself important for mRNA stability under purine stress [Martin, 2014], this sequence was included in the *LdNT2/NLuc* reporter construct to maintain endogenous-like expression. In parasites expressing *LdNT2/NLuc* flanked by wildtype UTRs, NLuc activity was robustly (~9.5-fold) upregulated after 48 hours of purine starvation (Figure 1B). As anticipated, cell lines harboring the 5’ sequence from *LdNT4* displayed an equivalent magnitude of NLuc induction, indicating that the *LdNT2* 5’-UTR is not required for purine-responsive control.

**Figure 1:**
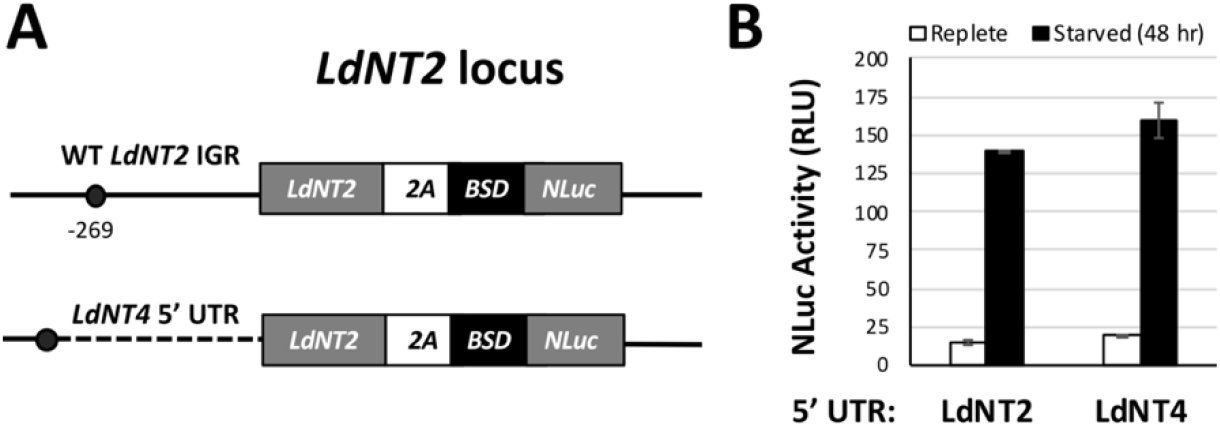
The LdNT2 mRNA 5’-UTR is not required for robust purine-responsive regulation. A) Top: Schematic of multicistronic constructs (referred to as *LdNT2/NLuc*) used to study *cis-* regulation of the *LdNT2* message. *NLuc* was fused via its N-terminus to the blasticidin resistance gene (*BSD*) to enable mutant selection. The *LdNT2* CDS was included to maintain endogenous-like levels of expression, followed by the self-cleaving *Thosea asigna* virus 2A peptide (2A). For control cell lines, unmodified up- and downstream *LdNT2* intergenic regions (IGRs) were used to direct construct integration into the endogenous *LdNT2* locus. In this configuration, both stability and translation of the *NLuc* message are governed by the native *LdNT2* UTRs and/or CDS. Co-translational 2A cleavage separates the post-translational fate of LdNT2 from that of the downstream BSD-NLuc polypeptide [de Felipe, 2006]. Thus, NLuc activity does not reflect changes in LdNT2 protein stability, which could potentially mask any regulation conferred by the UTRs. Circle represents the dominant *LdNT2* splice acceptor site, 269 nt upstream of the translation start. Bottom: The *LdNT2* 5’-UTR was replaced with that of *LdNT4* to verify that this region is not required for regulation under purine stress. Solid and dashed lines indicate purine-responsive and - unresponsive mRNA UTRs, respectively. Not pictured: The second allelic copy of *LdNT2* was replaced with a phleomycin resistance gene (Phleo). A firefly luciferase-puromycin resistance gene fusion (*Fluc-PAC)* integrated in place of the purine-unresponsive gene, UMP synthase (UMPS), was used as an internal control to normalize NLuc activity between replicates [Soysa, 2014]. B) Normalized NLuc activity from cell lines depicted in A, after 48 hours of culture in the presence or absence of purines. Figure shows the mean and standard deviation of experiments performed in biological duplicate.

We next performed serial deletions to identify the *cis*-acting elements encoded within the *LdNT2* 3’-UTR. As determined by 3’ Rapid Amplification of cDNA Ends (3’ RACE), the preferred *LdNT2* 3’ polyadenylation site lies 1.186 kb downstream of translation stop (Figure 2A). An *LdNT2/NLuc* reporter construct harboring wildtype *LdNT2* UTRs was therefore modified to tile this region with overlapping ~50-200 nt deletions and the effect on purine-responsive expression was examined. Deletions spanning the first 571 (Δ1 – Δ6) and last 511 (Δ7 – Δ11) bases had little to no effect on NLuc activity (Figure 2B). However, regulation was completely eliminated by deletion of 79 nts near the UTR midpoint. As this region was found to lie almost completely coincident with a 76 nt-long polypyrimidine tract, it is referred to as ΔCU.

**Figure 2:**
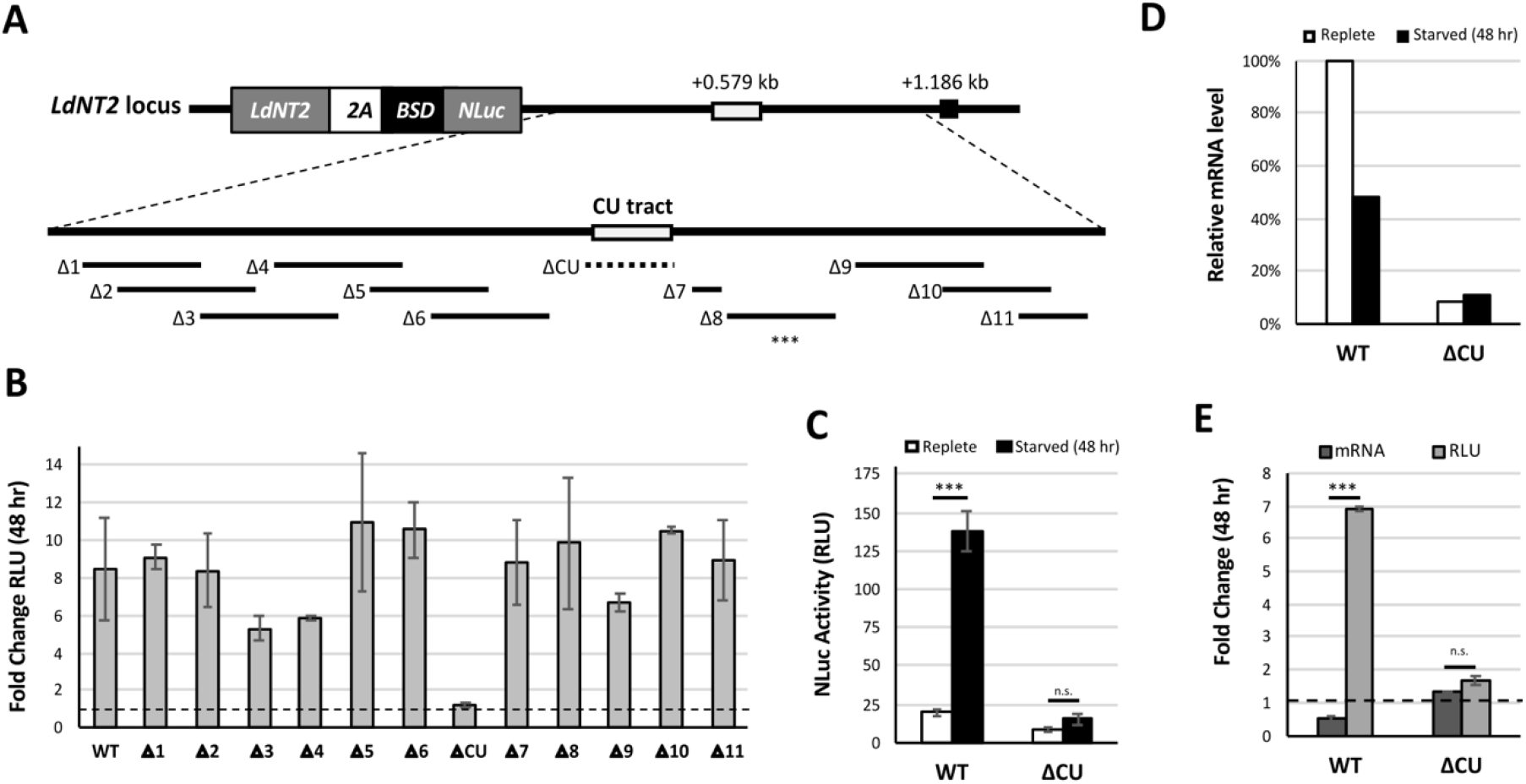
Deletional mutagenesis of the *LdNT2* 3’-UTR reveals that purine-responsive regulation is governed by a 76 nt-long polypyrimidine tract. A) Overlapping ~50-200 nt deletions (Δ1 -Δ11) generated in the 3’-UTR of the *LdNT2/NLuc* reporter construct, starting immediately downstream of translation stop and ending before the preferred polyadenylation site (black square). Dashed line highlights the region required for regulation, containing the *LdNT2* polypyrimidine (CU) tract. Numbers refer to distance between the indicated feature and translation stop. Not pictured: The second allelic copy of *LdNT2* was replaced with a phleomycin resistance gene (Phleo). A firefly luciferase-puromycin resistance gene fusion (*Fluc-PAC)* expressed from the *UMPS* locus was used as an internal control to normalize NLuc activity between replicates [Soysa, 2014]. B) Fold change in normalized NLuc activity after 48 hours of purine starvation, measured from cell lines depicted in A. WT refers to parasites expressing the *LdNT2/NLuc* construct under the control of native *LdNT2* UTRs. C-E) Investigating the contribution of the *LdNT2* polypyrimidine tract to mRNA stability and/or translation by paired dual-luciferase and RT-qPCR analyses. Where indicated, bars and error bars represent the mean and standard deviation of assays performed with 2 independent clones. C) Normalized NLuc activity measured after 48 hours of culture in the presence or absence of purines. D) Quantitation of *LdNT2/NLuc* transcripts from cultures assayed in C. Relative message level was determined by the comparative CT method using *UMPS* as an endogenous control gene and ‘WT replete’ as a reference sample. For ΔCT values and analysis details, refer to Table S1D. E) Fold change in *LdNT2/NLuc* mRNA level and NLuc activity after 48 hours of purine starvation. The comparative CT method was used to determine mRNA fold change for individual clones as shown in Table S1E. Single-factor ANOVA was calculated with Excel Descriptive Statistics Toolpak: \*\*\**P* ≤ 0.001; n.s., *P*>0.05.

Polypyrimidine tracts are ubiquitous in eukaryotic RNA. Through association with a variety of polypyrimidine tract binding proteins, including polypyrimidine tract binding protein 1 (PTB1), these features govern multiple stages of mRNA metabolism, including splicing, polyadenylation, nuclear export, and mRNA stability [reviewed in Romanelli, 2013]. The *T. brucei* PTB1 homolog, DRBD3, is an essential 37-kDa protein that binds mRNA at a conserved TTCCCCTCT motif [Das, 2015]. We observed that the *LdNT2* polypyrimidine tract encodes two overlapping, DRBD3-like binding sites (Figure 3A). Neither is perfectly identical to the published *T. brucei* DRBD3 consensus; however, in each, the identities of the divergent bases are among those tolerated for recognition [Das, 2015]. In bloodstream-form trypanosomes, RNAi knockdown of DRBD3 destabilized many differentially expressed transcripts, including the *LdNT2* ortholog, *TbNT10* [Estevez, 2008; Stern, 2009]. These observations strongly suggested DRBD3 as a potential candidate for interaction with the *LdNT2* polypyrimidine tract. We generated *LdNT2/NLuc* constructs lacking both putative DRBD3 motifs (ΔDRBD3) to test whether these regions were specifically required for regulation by the *LdNT2* 3’-UTR. Surprisingly, after 48 hours of purine stress, ΔDRBD3 transgenic parasites displayed an equivalent magnitude of NLuc induction to those expressing the same construct flanked by WT *LdNT2* UTRs (Figure 3B). Thus, regulation conferred by the *LdNT2* polypyrimidine tract cannot be attributed to this particular region.

**Figure 3:**
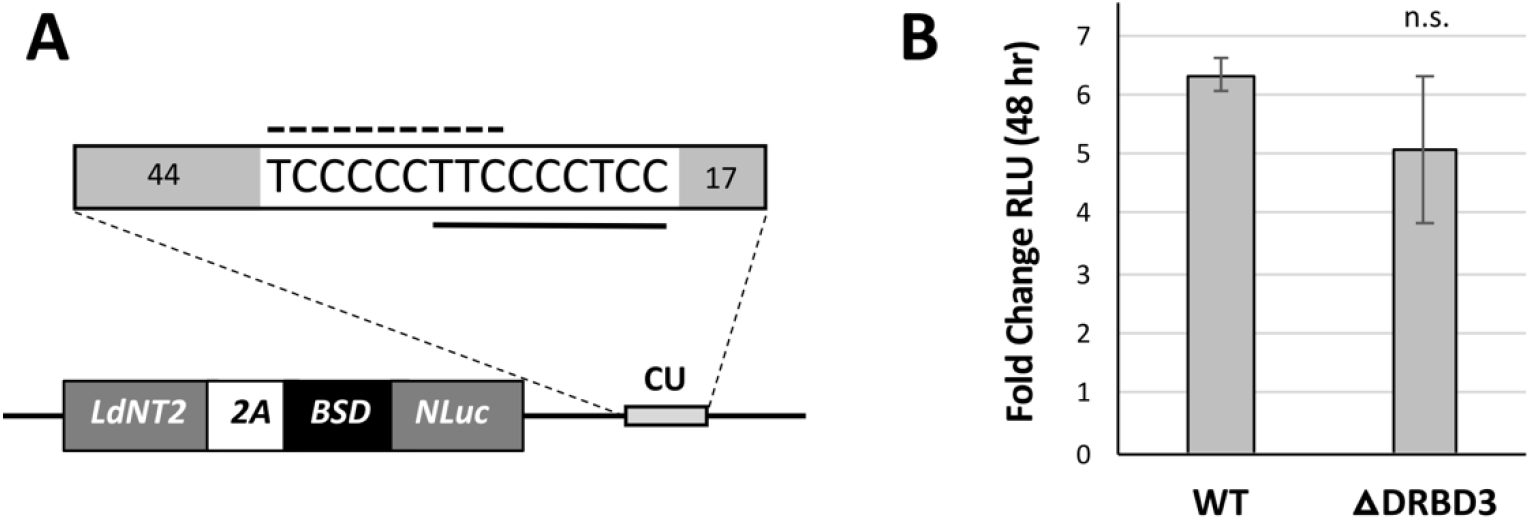
Binding sites for a known PTB protein in trypanosomes are encoded by the *LdNT2* polypyrimidine tract but are not required for regulation. A) Top: Sequence and relative position of two near-consensus DRBD3 binding sites in the *LdNT2* polypyrimidine tract (CU). Solid line indicates motif that differs from the published *T. brucei* DRBD3 consensus by a single residue; dashed line indicates a second, more degenerate site. In both cases, the divergent bases are tolerated for DRBD3 recognition [Das, 2015]. Numbers indicate the length of regions preceding and following putative DRBD3 motifs within the *LdNT2* polypyrimidine tract (76 nt in total). Diagram is not to scale. Bottom: *LdNT2/NLuc* reporter constructs were generated lacking putative DRBD3 motifs and integrated at the endogenous *LdNT2* locus. B) Regulation conferred by *LdNT2* mRNA UTRs, with and without DRBD3-like binding motifs. Data are normalized to *Fluc-PAC* expressed from the *UMPS* locus in the same cell line [Soysa, 2014]. Bars represent the mean and standard deviation of experiments performed in biological duplicate. Single-factor ANOVA was calculated with Excel Descriptive Statistics Toolpak. n.s., *P*>0.05.

### The polypyrimidine tract is a major determinant of LdNT2 transcript abundance and is required for translational enhancement under purine stress

*LdNT2* is unique in that it falls in the top 99.8^th^ percentile in mRNA abundance in log-stage *L. donovani* promastigotes [Martin, 2014]. At the same time, it ranks only in the 16^th^ percentile in terms of ribosome occupancy [Bifeld, 2018]. It is also known that while translation of the *LdNT2* message is significantly upregulated in response to purine starvation, mRNA levels do not change [Martin, 2014]. Based on these data, we hypothesized that *LdNT2* translation is somehow restricted or repressed under purine-replete conditions. To test whether the *LdNT2* polypyrimidine tract affects either mRNA abundance or translation with respect to purines, total RNA was isolated from the cultures analyzed in Figure 2C and *LdNT2/NLuc* transcripts were quantified via RT-qPCR (Figure 2D). For either cell line, the impact of purine starvation on *LdNT2/NLuc* mRNA level was then compared against the corresponding fold-change in NLuc activity to determine the translational contribution to regulation, reflected in the disparity between these two metrices (Figure 2E).

For parasites expressing *LdNT2/NLuc* flanked by wildtype UTRs, purine deprivation led to an approximate 50% decrease in transcript level. At the same time, NLuc activity was ~7-fold upregulated, suggesting that the *LdNT2* 3’-UTR and/or CDS together mediate a ~14-fold increase in translation under purine stress (Figure 2E, WT: dark vs light bars). This robust translational change is consistent with previous data for the endogenous *LdNT2* message [Martin, 2014]. Remarkably, *LdNT2/NLuc* mRNA abundance was reduced by 92% among parasites lacking the polypyrimidine tract (Figure 2D, white bars), implicating this sequence as an important determinant of message stability. At the protein level, the impact of polypyrimidine tract deletion was more modest, with ΔCU parasites demonstrating a 54% reduction in NLuc activity (Figure 2C, white bars). This disproportionate effect is in agreement with our hypothesis that a substantial portion of the *LdNT2/NLuc* mRNA pool is not translated under replete conditions. Furthermore, NLuc activity increased just ~1.7-fold in purine-starved ΔCU mutants, roughly equivalent to the change in mRNA abundance measured from the same cell line (Figure 2E, ΔCU: dark vs light bars). Thus, the translational component to regulation was completely eliminated by deletion of the *LdNT2* polypyrimidine tract. Taken together, the data support a model wherein the *LdNT2* message is poorly translated at steady-state but is maintained in high abundance by the polypyrimidine tract, such that it is available for translation under purine stress.

### The LdNT2/NLuc transcript localizes to discrete cytoplasmic foci under purine-replete conditions

We next considered potential mechanisms that could account for both the exceptional abundance of *LdNT2* mRNA and its low translational efficiency. One possibility is that *LdNT2* messages are stored in ribonucleoprotein (RNP) granules under purine-replete conditions, where they are simultaneously protected from degradation and sequestered away from the translational machinery. In this scenario, purine starvation is expected to trigger release of sequestered messages, making them available for translation. In addition, this model implies that the polypyrimidine tract stabilizes *LdNT2* mRNA by contributing to its sequestration. To test these possibilities, the distribution of transcripts, with or without the polypyrimidine tract, was examined via RNA fluorescence in situ hybridization (RNA-FISH) under purine replete and depleted conditions. The ‘WT’ and ΔCU reporter lines described in Figure 2 were cultured for 48 hours in the presence or absence of purines and *LdNT2/NLuc* mRNAs were visualized using fluorescent probes specific to the *LdNT2* CDS.

In parasites expressing *LdNT2/NLuc* under the control of wildtype *LdNT2* UTRs (Figure 4A, WT), staining was enriched in discrete foci under purine-replete conditions. This observation supports our hypothesis that the transcript is sequestered in replete cells. Further, although a punctate RNA-FISH signal was also detected in these parasites after 48 hours of purine starvation, the fluorescence intensity per cell was ~32% lower (Figure 4D and G, blue plots). Considering that aggregated transcripts are readily detected by RNA-FISH but individual molecules are not, this reduction in signal in purine-starved parasites could possibly reflect a decrease in compartmentalization of the *LdNT2/NLuc* message, consistent with release from storage for translation. In the case of *LdNT2/NLuc* transcripts lacking the polypyrimidine tract (Figure 3E, ΔCU), staining was also localized to discrete foci under purine stress. We suspect that, in both WT and ΔCU lines, these puncta may represent *LdNT2/NLuc* messages aggregated in nutrient stress granules, the formation of which is well-documented in purine-starved *Leishmania* [Shrivastava, 2019]. Interestingly, however, the RNA-FISH signal did not exceed background levels in the replete ΔCU sample (Figure 4B and G; red plot). Thus, deletion of the *LdNT2* polypyrimidine tract appeared to disrupt compartmentalization of *LdNT2/NLuc* mRNA but specifically under purine-replete conditions. Further experiments will be required to more thoroughly assess transcript dynamics into and out of sequestration and to understand the nature of granules formed under purine-replete and starvation conditions (see Discussion).

**Figure 4:**
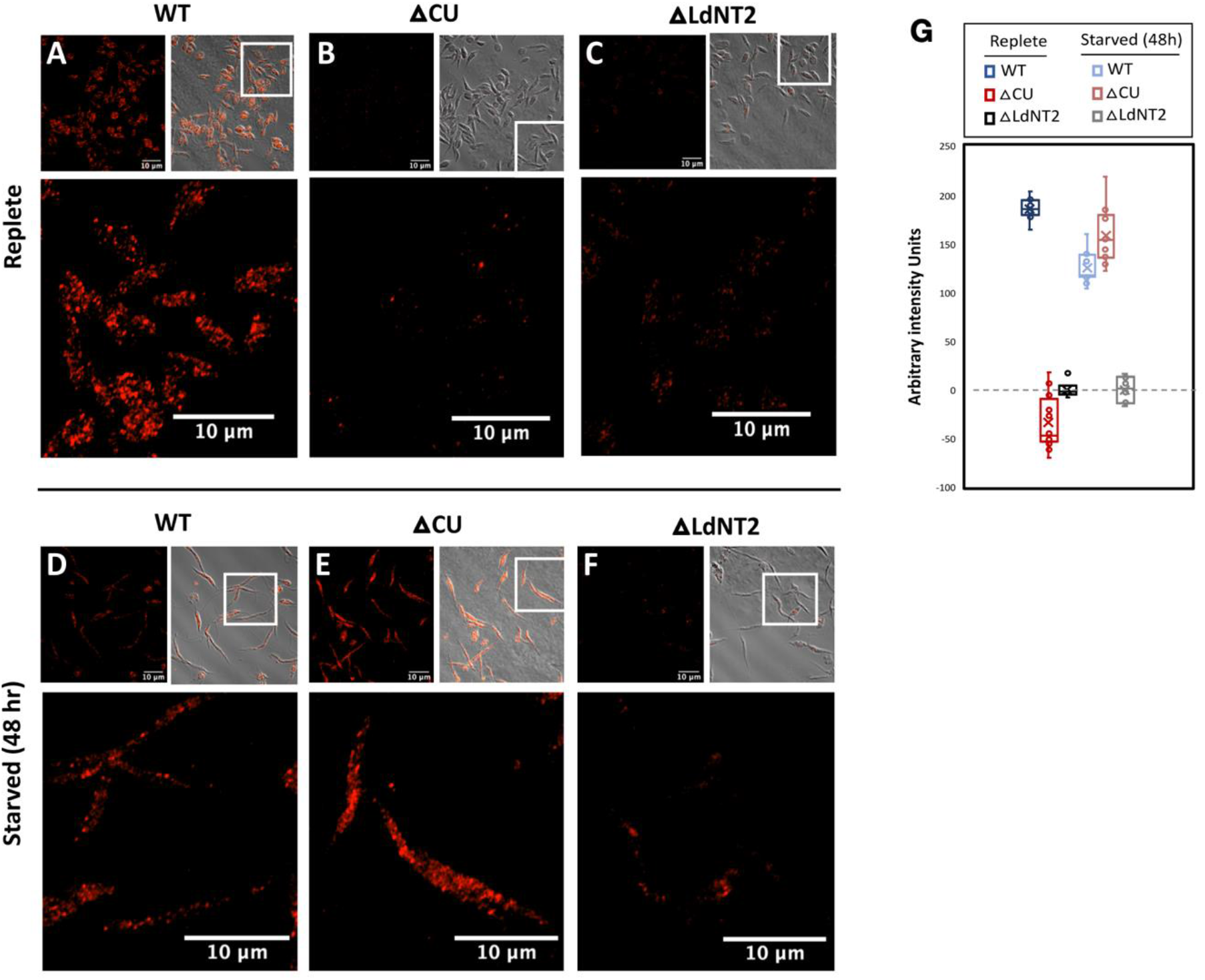
The *LdNT2* transcript localizes to discrete foci under purine-replete conditions. Promastigote parasites cultured for 48 hours in the presence (A-C) or absence (D-F) of purines were fixed on slides and processed for RNA-FISH. WT and ΔCU refer to the *LdNT2/NLuc* reporter lines of the same name depicted in Figure 2. *LdNT2/NLuc* transcripts were detected using fluorescently labeled probes specific for the *LdNT2* CDS. A homozygous *LdNT2* knockout line (ΔLdNT2) served as a control for nonspecific background. All fluorescent images were collected using the same settings; however, for qualitative assessment, representative images in A-C vs D-F were adjusted separately. [Author’s note: the elongated morphology visible in panels D-F is characteristic of purine-starved *Leishmania*. This phenomenon is well-documented and discussed elsewhere]. G) Graph reflects the average fluorescent intensity per pixel per cell measured from several representative fields (n=12 in Wildtype and ΔCU, n=6 in ΔLdNT2). Under either condition, data were corrected by subtracting the average of the intensity values collected in the ΔLdNT2 control sample.

### Purine-responsive regulation of the LdNT1 nucleoside transporter requires cooperation between two distinct cis-acting elements

In *L. donovani*, NT1 is encoded by two tandem, closely related genes: *LdNT1.1* and *LdNT1.2*. Both are functional when transfected into *Xenopus* oocytes; however, only *LdNT1.1* is expressed in promastigote parasites [Vasudevan, 1998]. For simplicity, all reference to *LdNT1* throughout pertains specifically to the gene expressed from the *LdNT1.1* locus.

By 3’ RACE, we determined that the *LdNT1* message is polyadenylated at two positions, yielding 3’-UTRs of either 1.681 or 1.819 kb in length (Figure 5A). Approximately 1 kb downstream of translation stop, we noted a 48 nt polypyrimidine tract, reminiscent of the purine-response element described for *LdNT2*. We therefore performed a focused molecular dissection of the surrounding sequence to test if this region also confers sensitivity to purines. A firefly luciferase-neomycin resistance gene fusion (*Fluc*-*NEO*) was expressed from the endogenous *LdNT1* locus under the control of either wildtype UTRs or a 3’-UTR harboring one of the ~50 nt deletions depicted in Figure 5A. Parasites were subjected to 48 hr of purine starvation and the impact on Fluc activity was evaluated. Luciferase induction was lost in all mutants lacking the *LdNT1* polypyrimidine tract (Δ6 -Δ7), consistent with the sequence functioning as a regulator of purine-responsive expression. However, several deletions preceding the polypyrimidine tract (Δ5, ΔUE1) also prevented regulation. To a resolution of 25 nt (i.e. the length of overlap between adjacent deletions), we determined that the 5’ boundary of the *LdNT1* regulator lies 46 nt upstream of the polypyrimidine tract (position +0.910). The intervening sequence is heretofore referred to as the *LdNT1* upstream element, or UE1.

**Figure 5:**
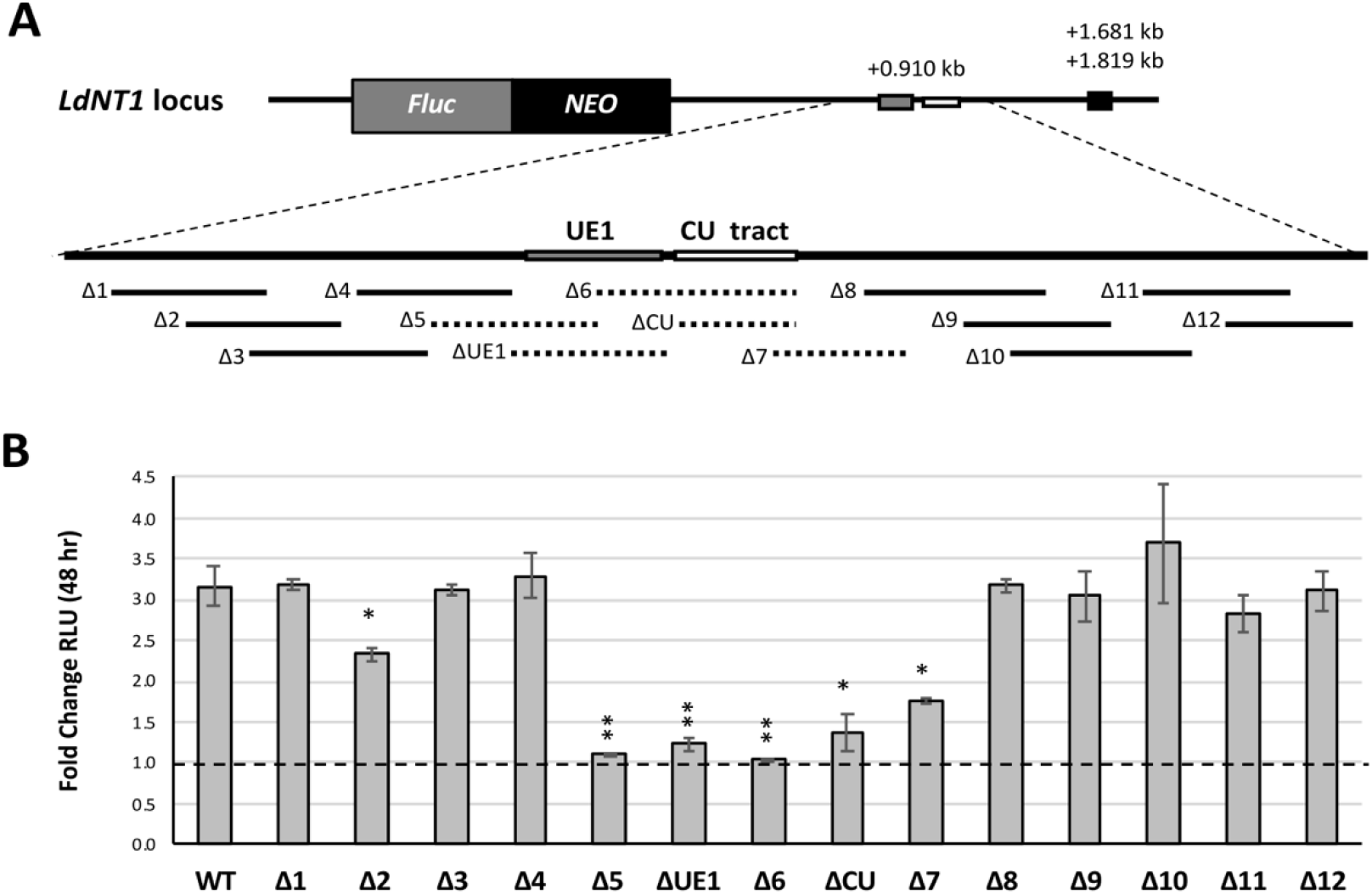
Deletional mutagenesis of the *LdNT1* 3’-UTR reveals a bipartite purine-response element. A) Firefly luciferase (*Fluc*) was fused in-frame to a selectable neomycin resistance marker (*NEO*) and flanked with the *LdNT1* IGRs to direct construct integration into the endogenous locus. Overlapping deletions (Δ1 - Δ12) encompass ~50 nt each, overlap by ~25 nt apiece, and span a total distance of 420 nt around the *LdNT1* polypyrimidine (CU) tract. Dashed lines indicate regions required for purine-responsive regulation, corresponding to the CU tract and upstream element (UE1). Black square represents the dominant *LdNT1* polyadenylation site(s) as determined by 3’-RACE. Numbers refer to distance between the indicated feature and translation stop. Not pictured: A *Renilla* luciferase-puromycin resistance gene fusion (*Rluc-PAC*) expressed from the *UMPS* locus serves as an internal normalization control. B) Fold change in normalized Fluc activity after 48 hours of purine starvation, measured from cell lines depicted in A. WT refers to parasites expressing *Fluc-NEO* under the control of native *LdNT1* UTRs. Bars represent the mean and standard deviation of experiments performed in biological duplicate. Single-factor ANOVA was calculated with Excel Descriptive Statistics Toolpak: \**P* ≤ 0.05, \*\**P* ≤ 0.01.

It is known that increases in both mRNA abundance and translational efficiency contribute to LdNT1 upregulation under purine stress [Martin, 2014]. We therefore asked if either of these processes are affected by the *LdNT1* polypyrimidine tract and/or UE1. The WT, ΔCU, and ΔUE1 reporter lines analyzed in Figure 5 (represented separately in Figure 6A for clarity) were cultured with and without purines for 48 hours. As described previously, the relative contributions of mRNA stability versus translation were then determined by comparing the purine-responsive change in *Fluc-NEO* mRNA abundance against the corresponding change in luciferase activity for each cell line. In the absence of purines, transgenic parasites harboring wildtype *LdNT1* UTRs demonstrated a 30% reduction in *Fluc-NEO* mRNA (Figure 6B, WT). This is in contrast to what has been reported for the endogenous *LdNT1* message, which is modestly but significantly upregulated in purine-starved *L. donovani* promastigotes [Martin, 2014]. We interpret this to mean that the *LdNT1* CDS contributes to mRNA stability under purine starvation. Nonetheless, luciferase activity increased ~2.5-fold in the same cell line, pointing to a ~3.5-fold increase in translation mediated by the *LdNT1* UTRs (Figure 6B, WT: disparity between dark vs light bars). Interestingly, deletion of either the *LdNT1* polypyrimidine tract or UE1 had differing effects on abundance and translation of the reporter construct. In purine-starved parasites lacking the polypyrimidine tract, for instance, *Fluc-NEO* transcripts were even more substantially decreased than in the WT reporter line (73% vs 30%, respectively), suggesting that this region confers stability to the *LdNT1* message under purine stress. Yet these cells still displayed a robust 4.8-fold increase in translation (Figure 6B, ΔCU: dark vs light bars). In contrast, deletion of UE1 completely eliminated the translational contribution to regulation but did not negatively impact mRNA level (Figure 6B, ΔUE1). Taken together, these data suggest that the *LdNT1* polypyrimidine tract and UE1 function independently of each other to confer regulation at the levels of mRNA abundance and translation, respectively.

**Figure 6:**
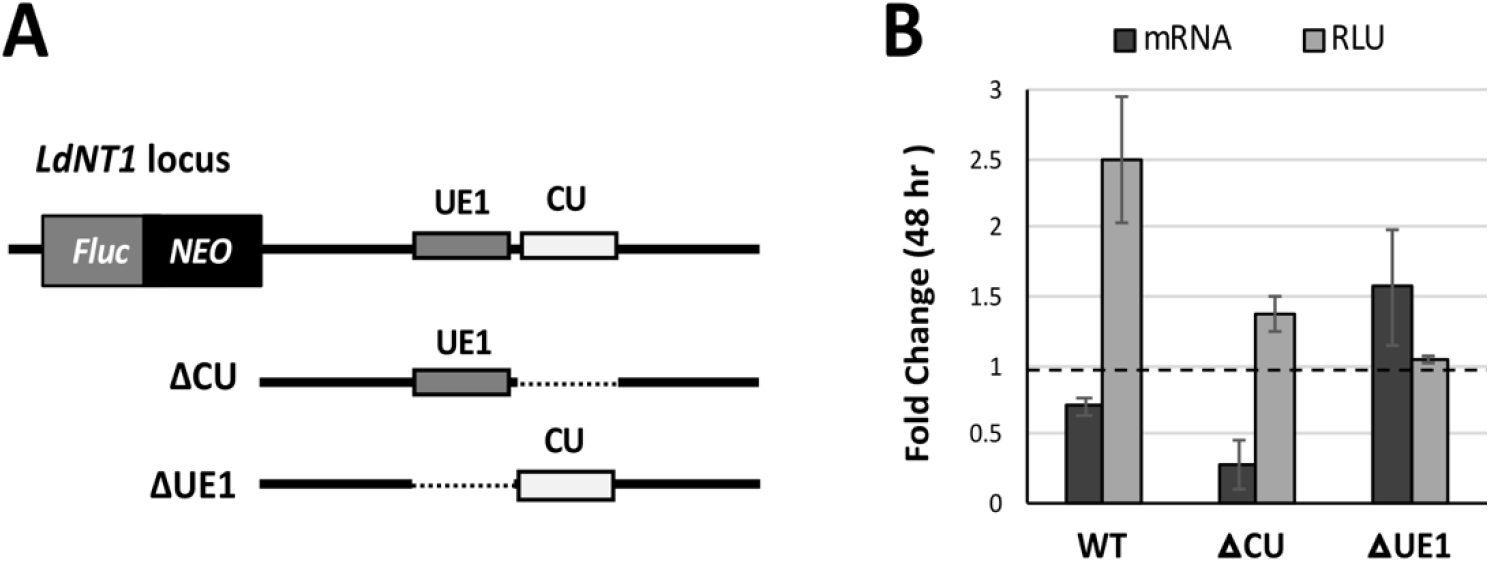
The *LdNT1* upstream element and polypyrimidine tract govern translation and mRNA stability independently. A) Investigating the contributions of the *LdNT1* upstream element (UE1) and polypyrimidine tract (CU) to mRNA stability and/or translation by paired dual-luciferase and RT-qPCR analyses. *Fluc-NEO* flanked by wildtype *LdNT1* UTRs serves as a control (referred to as WT in text). ΔCU and ΔUE1 cells lines are the same as those of the same name in Figure 5A but for ease of interpretation, their 3’-UTRs are represented again here. Not pictured: *Rluc-PAC* expressed from the *UMPS* locus was used to normalized between experiments. B) Fold change in *Fluc-NEO* mRNA level and Fluc activity after 48 hours of purine starvation, measured from the cell lines depicted in A. The mRNA fold change for individual clones was determined by the comparative CT method as shown in Table S2. Bars represent the mean and standard deviation of experiments performed in biological duplicate.

### The LdNT1 polypyrimidine tract confers regulation in the context of the LdNT2 mRNA 3’-UTR, in the absence of UE1

For both *LdNT1* and *LdNT2*, purine-responsive expression is governed by a polypyrimidine tract in the mRNA 3’-UTR. However, in the latter case, regulation also requires cooperation with an adjacent translational enhancer (i.e. UE1). In a final experiment, we asked whether just the polypyrimidine tract from *LdNT1* could substitute for that of *LdNT2* to confer regulation outside of its endogenous genetic context. We swapped these respective elements in the *LdNT2/NLuc* reporter construct and evaluated the impact on NLuc expression after 48 hours of purine stress. As depicted in Figure 7, *LdNT2/NLuc* flanked by wildtype *LdNT2* UTRs (WT) displayed a ~7.7-fold increase in luciferase activity. Remarkably, reporter lines harboring a polypyrimidine tract from *LdNT1* not only retained purine-responsive expression, but the magnitude of induction was ~7-fold greater than regulation conferred by the endogenous sequence (Figure 7, LdNT1). This suggests that, at least in the particular genetic context of the *LdNT2/NLuc* construct, the *LdNT1* polypyrimidine tract is sufficient to confer purine-sensitivity independently of UE1.

**Figure 7:**
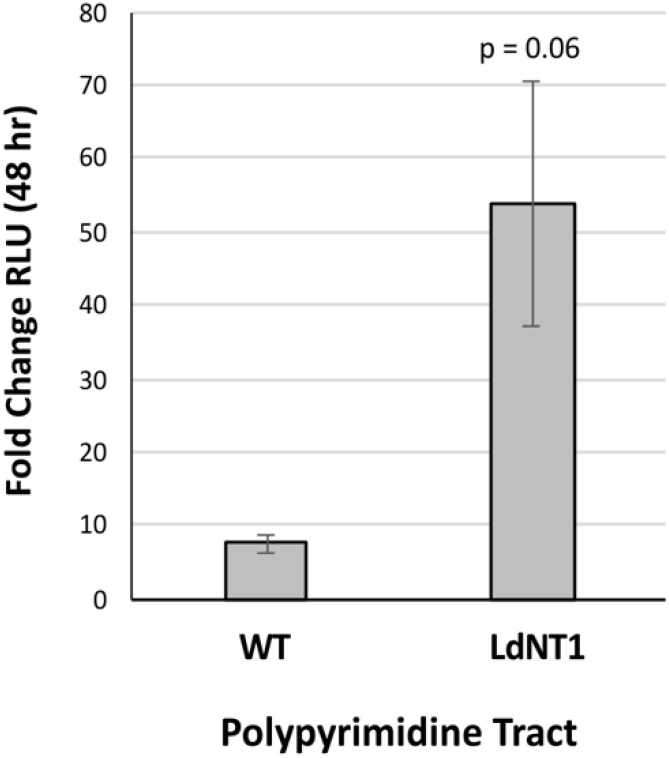
The *LdNT1* polypyrimidine tract confers purine-responsive regulation when substituted at the *LdNT2* locus. In the *LdNT2/NLuc* construct (depicted in Figure 1A, top), the *LdNT2* polypyrimidine tract was replaced with that of *LdNT1*. WT refers to *LdNT2/NLuc* flanked by wildtype *LdNT2* 5’- and 3’-UTRs. Graph displays fold change in NLuc activity after 48 hours of purine starvation. NLuc activity is normalized to Fluc expressed from the endogenous UMPS locus. Bars represent the mean and standard deviation of experiments performed in biological duplicate.

## Discussion

We have used the purine transporters as a model to examine regulation of the *Leishmania* purine stress response. Previous studies suggested that these genes are governed by different post-transcriptional mechanisms and we recently described a repressive stem-loop in the 3’-UTR of *LdNT3* that serves to limit expression under purine-replete conditions [Licon, 2020]. Though conserved in orthologous genes from a variety of kinetoplastids, the *LdNT3* stem-loop was not found elsewhere in the *L. donovani* genome, including the other LdNTs. Thus, in the present work, we performed systematic deletion analysis of the *LdNT1* and *LdNT2* UTRs to identify the elements responsible for their control.

*LdNT2* mRNA is exceptionally abundant yet poorly translated in purine-replete *L. donovani* promastigotes. We found that regulation depends on 76 nt-long polypyrimidine tract, encoded in the mRNA 3’-UTR. In the context of a reporter construct, we showed that mRNA abundance is ~92% reduced by deletion of the *LdNT2* polypyrimidine tract (Figure 2D). Translational enhancement under purine stress was also eliminated (Figure 2E). Based on these observations, we suggest a model for *LdNT2* regulation wherein i) in replete cells, *LdNT2* messages are simultaneously stabilized and sequestered away from the translational machinery by storage in RNP granules, ii) translation is upregulated when stored transcripts are made accessible by trafficking out of granules in response to purine stress, and iii) *LdNT2* mRNA stability and sequestration are dependent on the polypyrimidine tract. Although the results of our RNA-FISH analysis are not entirely conclusive, they provide compelling preliminary evidence in support of this model.

Under purine-replete conditions, we found that transcripts harboring wildtype *LdNT2* UTRs localize to discrete, cytoplasmic foci (Figure 4A). This observation is generally in line with our model. However, it begs the question: what is the nature of the RNA-containing structures? Many types of RNP granules have been described in kinetoplastid parasites, with distinct protein markers identified. Broadly speaking, these are distinguished based on whether they form under favorable or stress conditions, and whether transcripts are stored or degraded within [reviewed Kramer, 2014]. For example, both heat shock and nutrient restriction induce formation of cytoplasmic stress granules, which act to store and protect mRNAs until stress is resolved [Kramer, 2008; Fritz, 2015]. Specialized sites of mRNA turnover known as processing bodies (P-bodies) have also been observed. As in other eukaryotes, P-bodies are constitutively present in kinetoplastids but increase in size and abundance in response to environmental insults [Holetz, 2007; Kramer, 2008, Fritz, 2015]. Recently, a paralog of eukaryotic initiation factor eIF4E was found to concentrate in cytoplasmic granules in purine-starved *Leishmania* promastigotes. These structures also contained ribosomal subunits and mature mRNAs, suggestive of a role in translational repression [Shrivastava, 2019]. We suspect that the fluorescent puncta observed in purine-starved *LdNT2/NLuc* parasites (Figure 4D and E) represent one or more of these established granule types. However, to our knowledge, there have been no examples of translational repression by selective mRNA sequestration described in any kinetoplastid parasite to date, making the phenomenon we have described in purine-replete cells potentially novel. Future efforts to characterize the protein composition of these granules may be informative in discerning their true function with respect to the *LdNT2* transcript.

In the same ‘WT’ cell line, intensity of the RNA-FISH signal was significantly reduced after exposure to purine stress (Figure 4D and G). As aggregated RNAs are more readily detected by RNA-FISH than are individual molecules, this could reflect a general shift in the *LdNT2/NLuc* mRNA pool from a sequestered to a free state. Such a phenomenon is consistent with our hypothesis that *LdNT2* transcripts are trafficked out of storage granules to be translated in response to purine starvation. However, because the overall *LdNT2/NLuc* message level is also reduced under stress (Figure 2D, WT), we are cautious in accepting this interpretation. As more conclusive evidence, single molecule RNA-FISH (smRNA-FISH) could provide insight into the dynamics of individual transcripts, into and/out of sequestration. It could also potentially be informative to compare the distribution of *LdNT2* to other messages for which purine-responsive localization has already been determined. *HSP83*, for example, yields a diffuse cytoplasmic signal in replete *Leishmania* and accumulates in storage granules under purine restriction [Shrivastava, 2019]. In any case, further examination is required to verify the significance of this observation.

Our preferred model states that storage of the *LdNT2* message in RNP granules physically protects it from the degradation machinery in the cytosol. In this scenario, message stability is dependent upon sequestration. Working backward from our observation that the *LdNT2* polypyrimidine tract is a major regulator of mRNA abundance (Figure 2D), one would therefore predict that transcripts lacking this element are not recruited to granules under purine-replete conditions. Alternatively, it is also possible that *LdNT2* mRNA abundance is controlled independently of sequestration, with the polypyrimidine tract only influencing the former. In this case, *LdNT2* transcripts would be expected to accumulate in cytoplasmic foci in replete cells, irrespective of the polypyrimidine tract. In purine-replete ΔCU *LdNT2/NLuc* parasites, we did not readily detect fluorescent puncta and the RNA-FISH signal did not exceed background levels, as measured from the ΔLdNT2 control (Figures 4B and G). One interpretation of these data is that ΔCU *LdNT2/NLuc* transcripts are not sequestered in granules, in line with our preferred model of regulation. However, because abundance of the *LdNT2/NLuc* message is also dramatically reduced in this cell line (~92% lower than WT; Figure 2D), it is equally possible that an absence of signal in these samples merely reflects a transcript that has dropped below the limit of detection. As evidence in support of the former case, fluorescent puncta were clearly visible in ΔCU parasites exposed to purine stress (Figure 4E), despite there being no significant difference at the mRNA level (Figure 2D, ΔCU). This observation would seem to suggest that *LdNT2/NLuc* mRNAs in the ΔCU cell line are not so low abundance as to be undetectable when aggregated. Thus, although the results of this experiment are not conclusive, we are encouraged that they genuinely reflect an inability of purine-replete *Leishmania* to compartmentalize transcripts lacking the *LdNT2* polypyrimidine tract. In future studies, smRNA-FISH will be conducted to distinguish between the two options outlined above.

As an important caveat, it should be noted that all of the observations described here are based on RNA-FISH experiments performed with a non-native RNA construct. *LdNT2/NLuc* encodes all components of the mature *LdNT2* message (i.e. 5’ and 3’ UTRs, CDS). In addition, several preliminary RNA-FISH experiments were conducted to detect the endogenous *LdNT2* transcript. Qualitatively, the subcellular distribution of this message appeared consistent with what we have reported for the ‘WT’ *LdNT2/NLuc* construct [data not shown]. However, we cannot exclude the possibility that aspects of regulation are affected. In future experiments, we will also verify the localizations described here with endogenous *LdNT2* mRNA.

We identified a second purine-responsive polypyrimidine tract in the *LdNT1* 3’-UTR. Like that of *LdNT2*, loss of this region had a negative impact on mRNA abundance. However, we found that translation of the *LdNT1* message is separately controlled by an adjacent sequence, termed UE1 (Figure 6). Both features are required for induction under purine stress, suggesting that they function cooperatively to enact changes in protein abundance. We were therefore surprised to find that, when substituted into the *LdNT2/NLuc* reporter construct, the polypyrimidine tract from *LdNT1* was sufficient for regulation, independent of UE1 (Figure 7). This observation speaks to the importance of local genetic context in *cis*-regulation, the subtleties of which are often overlooked. Indeed, the effect of substitution was robust, with the magnitude of reporter induction far exceeding that conferred by the wildtype *LdNT2* 3’-UTR under purine stress. The reason for this is unclear, particularly without knowing the impact of the substitution at the mRNA level. Does the *LdNT1* polypyrimidine tract confer greater message stability in this context, resulting in a larger pool of *LdNT2/NLuc* for translation under stress? By that same token, do *LdNT2/NLuc* messages harboring the *LdNT1* polypyrimidine tract also localize to cytoplasmic foci under replete conditions? The answers to these and other questions could provide further mechanistic insight into the respective functions of the two polypyrimidine tracts controlling nucleoside transport.

In combination with our previous work on LdNT3, we can now definitively state that each *L. donovani* purine transporter is controlled by distinct purine-response elements. However, the obvious question remains: What binds to these regions *in vivo?* We indirectly tested one candidate for *LdNT2* in the form of DRBD3. Despite strong evidence suggesting that this protein binds to and stabilizes transcripts encoding the *LdNT2* ortholog in *T. brucei* [Das, 2015; Estévez, 2008; Stern, 2009], deletion of two near-consensus DRBD3-binding sites in the *LdNT2* polypyrimidine tract had no significant effect on regulation. Admittedly, based purely on these results of this experiment, we cannot exclude the possibility of an interaction between *L. donovani* DRBD3 and *LdNT2*, occurring elsewhere in the *LdNT2* message or at the deleted sites. Further, as we did not examine the impact of this deletion at the transcript level, we cannot speculate as to a potential role for DRBD3 in regulating *LdNT2* mRNA abundance. We can merely state that the 15 nts deleted are not required for *LdNT2* induction in response to purines. In any case, given the mechanistic differences in regulation conferred by each of the *LdNT* purine-response elements (i.e. positive versus negative control, mRNA stability versus translation), we suspect that the protein factors involved differ between the transporters. Their characterization highlights a remarkable degree of complexity in the regulation of the *Leishmania* purine stress response.

## Materials and Methods

### Leishmania donovani culture

All cell lines described in this work were generated from the *L. donovani* 1S-2D clonal subline LdBob, originally provided by Dr. Stephen Bevereley [Goyard, 2003]. Promastigotes parasites were routinely cultured at 26 °C with 5% CO_2_ in Dulbecco’s Modified Eagle-Leishmania (DME-L) medium, supplemented with 5% SerumPlus^TM^ (SAFC BioSciences/Sigma Aldrich, St. Louis, MO; a purine-free alternative to standard FBS), 1mM L-glutamine, 1x RPMI vitamin mix, 10uM folic acid, 50 ug/ml hemin, and 100 uM hypoxanthine as a purine source. For culture maintenance, blasticidin, puromycin, and phleomycin were used at concentrations of 30 ug/ml, 25 ug/ml, and 50 ug/ml, respectively. For neomycin-resistant lines, G418 was added at 25 ug/ml. To elicit purine starvation, log-stage cultures were pelleted via centrifugation (5000 × g for 5 min), washed once in purine-free DME-L, and resuspended at a density of 1 × 10^6^ cells/ml in media lacking purines but containing all other nutrient supplements.

### Luciferase reporter constructs and cloning

With the exception of plasmid S1B (see below), targeting vectors were generated using the multi-fragment ligation approach described in [Fulwiler, 2011] and depicted in Figure S1A. For *LdNT2/NLuc* constructs, the *2A-BSD-NLuc* transgene fusion was provided by donor plasmid pCRm-2A-coBSD-NLuc [Yates, P., manuscript in preparation]. Targeting sequences used to direct integration into the *LdNT2* locus were PCR amplified from genomic DNA with Phusion High Fidelity DNA polymerase (New England Biosciences, Ipswitch, MA) using the primers listed in Table S3. All vector components were digested with SfiI, gel-purified, and assembled in a single ligation step.

The *LdNT2* 5’-UTR replacement construct depicted in Figure S1B was assembled in a step-wise fashion. First, genomic DNA was isolated from transgenic parasites expressing *LdNT2/NLuc* flanked by endogenous *LdNT2* intergenic regions (IGRs). Using the primers listed in Table S4, the entire *LdNT2/NLuc* reporter locus (including the preceding 5’-IGR) was PCR amplified and cloned into the SwaI restriction sites of plasmid pBB [Fulwiler, 2011]. The resultant vector (not pictured) was then subjected to whole plasmid amplification to exclude the 269 nt immediately upstream of translation start (i.e. the *LdNT2* 5’-UTR) and PCR products were DpnI-treated to eliminate template plasmid. A length of 250 nt immediately preceding *LdNT4* translation start was amplified from genomic DNA and both components were assembled via the Gibson Assembly method using NEBuilder HiFi DNA Assembly Master Mix (New England BioLabs, Ipswitch, MA).

For deletional mutagenesis of the *LdNT2* 3’-UTR, the primers listed in Table S5 were used to perform whole plasmid amplification of vector S1A, with each divergent primer pair excluding a ~50-200 nt-long region. PCR products were circularized via NotI restriction sites encoded in primer 3’ ends. A similar approach was taken to delete putative DRBD3 binding motifs from the *LdNT2* polypyrimidine tract, using the primers listed in Table S6. However, in this case, circularization was achieved by Gibson Assembly.

To introduce deletions into the *LdNT1* 3’-UTR, the downstream IGR was PCR amplified as two separate halves, with each separated by the intended deletion site. The selectable *Fluc-NEO* fusion (Figure S1C) was donated by vector pCRm-luc2-NEO (Genbank Accession number KF035120.1). Individual fragments were assembled using the multi-fragment ligation scheme shown in Figure S1A. Primers are listed in Table S7.

To substitute the *LdNT1* polypyrimidine tract for that of *LdNT2* in *LdNT2/NLuc* reporter constructs, XbaI restriction sites were introduced in place of the *LdNT2* polypyrimidine tract via whole plasmid amplification. The *LdNT1* polypyrimidine tract was then amplified from genomic DNA and inserted via directional XbaI cloning. All relevant primers are listed in Table S8.

### Transfections

In all luciferase assays described throughout work, a compatible luciferase expressed under the control of purine-unresponsive UTRs from the gene encoding UMP synthase (*UMPS*) served as a control from normalization. Hence, all Fluc- and NLuc-based reporter constructs were transfected into LdBob derivatives expressing either *Rluc-PAC* or *Fluc-PAC*, respectively, from the endogenous *UMPS* locus [Soysa, 2014]. Recipients of all *LdNT2/NLuc* constructs also harbored a heterozygous *LdNT2* deletion (*UMPS/umps::Fluc-PAC; LdNT2/ldnt2::Phleo)* such that, in the resultant cell lines, the only expressed copy of *LdNT2* was encoded by the reporter locus. For deletion mutagenesis of the *LdNT1* 3’-UTR, the *Fluc-NEO* reporter constructs depicted in Figure S1C were delivered to a recipient line expressing *Fluc-BSD* from the endogenous *LdNT1* locus (*UMPS/umps::Rluc-PAC; LdNT1/umps::Fluc-BSD*) such that integration was directed by the *Fluc* reporter gene.

All transfections were performed using at least 3ug SwaI-linearized plasmid DNA and following the high-voltage electroporation protocol described previously [Robinson, 2003]. Electroporated cells were transferred immediately into 10 mL of complete DME-L. To derive independent clones, 200 ul was then added to the first column of wells on a 96 well plate and cultures were subjected to 2-fold serial dilution. Transfectants were allowed to rest overnight at 26 °C in 5% CO_2_ before selection was initiated by the addition of 100 ul/well of 2X blasticidin (60 ug/ml) or, for *NEO*-encoding constructs, 2X G418 (50 ug/ml). Proper construct integration and, when applicable, inclusion of deletions, was PCR-verified for all clones.

### Dual-Luciferase analysis

Dual-luciferase assays were performed using either the Dual-Glo Luciferase Assay System (Fluc and Rluc) or the Nano-Glo Dual-Luciferase Reporter Assay (Nluc and Fluc) from Promega (Madison, WI). Analyses were performed on 35 ul of culture in white polystyrene 96-well half-area plates (Corning, Amsterdam). As described in the respective product technical manuals, plates were protected from light and shaken on an orbital shaker at room temperature (RT) for each 10-minute incubation step. Luminescence was measured using a Veritas Microplate Luminometer (Turner BioSystems, Sunnyvale, CA).

### RNA, cDNA, and RT-qPCR analysis

Total RNA was isolated from 5 × 10^6^ log-stage parasites with the NEB Monarch Total RNA miniprep kit and contaminating genomic DNA was eliminated with the TURBO DNA-*free* kit from ThermoFisher. Samples were then subjected to first-strand cDNA synthesis using the Applied Biosystems High Capacity cDNA Reverse Transcription Kit. RT-qPCR was performed using previously validated primers (Table S9; Martin, 2014) with 12 ng of cDNA and the NEB Luna Universal qPCR Master Mix. Reactions were run on an Applied Biosystems StepOnePlus instrument using the “Fast” ramp speed and the following thermocycling parameters: 95 °C for 60 seconds; 40 cycles of denaturing at 95 °C for 15 seconds followed by a 30 second extension at 60 °C. A final melt curve step was included to verify the specificity of amplification. Relative message abundance was determined using the comparative CT (ΔΔCT) method as described previously [Martin, 2014].

### RNA-FISH

To generate probes for RNA-FISH, the *LdNT2* CDS was amplified from genomic DNA using the ‘LdNT2 CDS’ primers listed in Table S3 and purified using the NEB Monarch PCR and DNA Cleanup kit. DNA probes were labeled for 4 hr via nick translation (Vysis; Abbott Laboratories, Abbott Park, Illinois) with Spectrum Orange-dUTP (Vysis) according to manufacturer instructions. Labeled probes were suspended in SLI/WCP hybridization buffer (Vysis) to a final concentration of 16.6 ng/ul.

For analysis of *L. donovani* promastigotes, parasites were pelleted by centrifugation and resuspended at a density of ~2×10^7^ cell/ml in 3.7% formaldehyde. To minimize clumping, cell pellets were gently disrupted by running the collection tube 5x along a microtube rack prior to fixation. Fixed cells (500 ul) were distributed over polylysine-coated slides and allowed to settle for ~20 minutes at RT before the cell suspension was aspirated from the slide surface. Slides were then stored in 70% EtOH at −20°C until use. For hybridization, labeled probes were denatured at 75 °C for 10 min. Just prior to probe application, slides were dehydrated in an EtOH series (3 min each in 90% and 100% EtOH) and allowed to air dry at RT. Slides were then hybridized with 20 ul of denatured probes in a humid chamber for 14-16 hours at 37°C. As an additional precaution against desiccation during hybridization, coverslip-mounted slides were also sealed with rubber cement. Post-hybridization washes consisted of i) one 3-minute wash in 2xSSC/50% formamide at 37°C and ii) one 1-minute wash in 2xSSC/0.1% Triton X-100 at RT. Coverslips were then mounted with Prolong Gold DAPI antifade and imaged on a ZEISS LSM 980 in Airyscan SR mode with a 63×1.4 NA objective.

For individual images, average fluorescence intensity per cell was quantitated as the signal intensity per pixel scaled to the number of parasites per field (counted by DAPI-stained kinetoplasts). To account for background, under either purine-treatment condition, the average intensity measured in the ΔLdNT2 control was subtracted from measurements collected under the same conditions in experimental cell lines. Thus, data are represented as delta values.

