## Supplemental Figure 1 for "Distinct *cis*-acting elements govern purine-responsive regulation of the *Leishmania donovani* nucleoside transporters, LdNT1 and LdNT2"

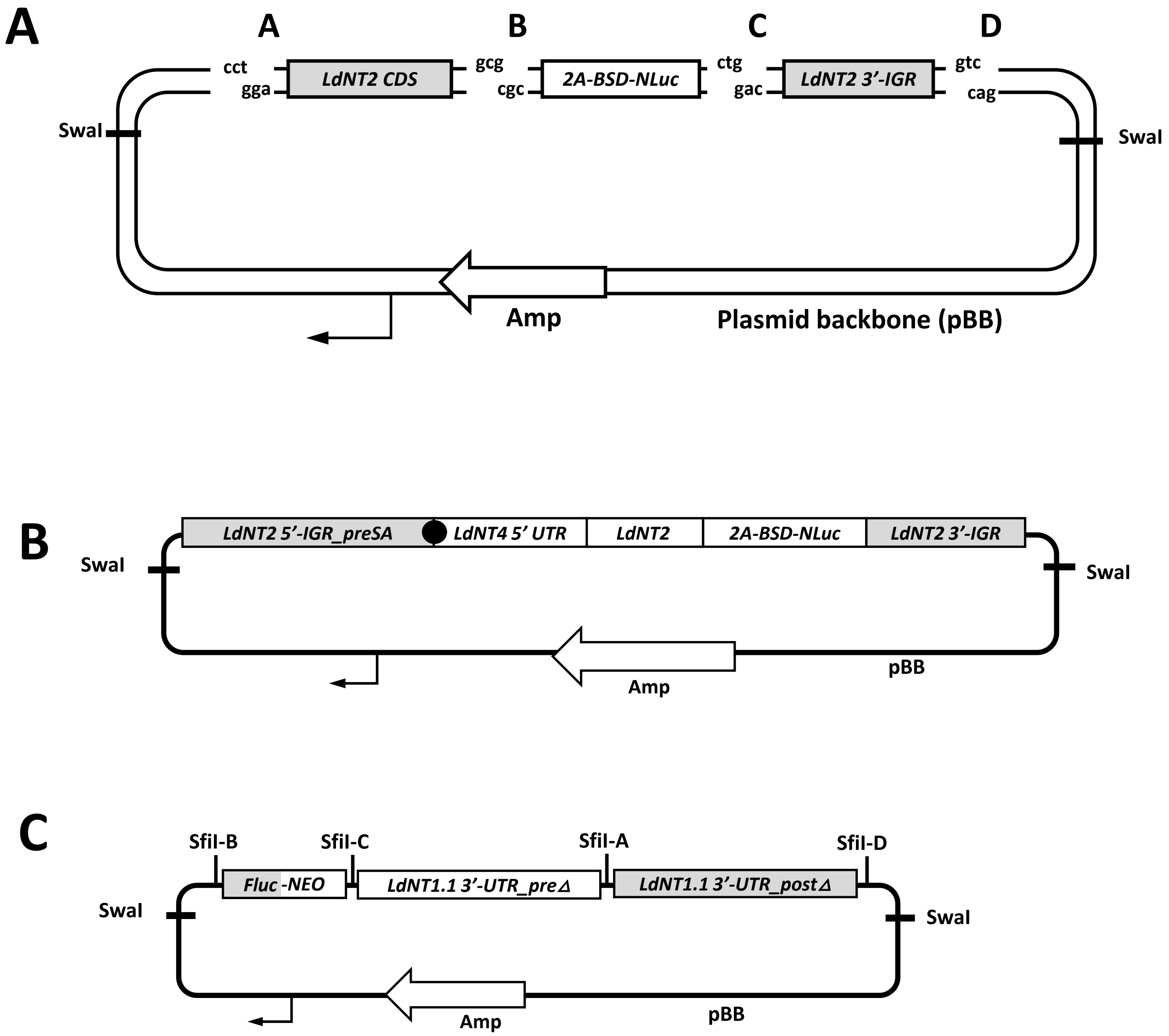
A
A
B
C
D
cct
gga
LdNT2 CDS
gcg
cgc
2A-BSD-NLuc
ctg
gac
LdNT2 3'-IGR
gtc
cag
SwaI
SwaI
Amp
Plasmid backbone (pBB)
B
LdNT2 5'-IGR_preSA
LdNT4 5' UTR
LdNT2
2A-BSD-NLuc
LdNT2 3'-IGR
SwaI
SwaI
Amp
pBB
C
SfiI-B
SfiI-C
SfiI-A
SfiI-D
Fluc-NEO
LdNT1.1 3'-UTR_preΔ
LdNT1.1 3'-UTR_postΔ
SwaI
SwaI
Amp
pBB

**Figure S1. Gene targeting vector assembly.** For simplicity, only the strategy for construction of the standard *LdNT2/NLuc* construct is shown here; however, all *LdNT1* 3'-UTR deletion mutagenesis constructs (C) were assembled using the same multi-fragment ligation approach. In all vectors, sequences used to direct targeted integration are shaded grey. A) Assembly of the *LdNT2/NLuc* reporter construct. Individual vector components are flanked by sites for restriction endonuclease SfiI. Restriction sites are designed to generate nonidentical 3'-overhangs upon digestion (designated as junctions A through D), facilitating ordered and directional plasmid assembly in a single ligation reaction. The minimal plasmid backbone (pBB) encodes an origin of replication (black arrow) and an ampicillin resistance gene (AMP). B) *LdNT2/NLuc* 5'-UTR replacement construct. The complete *LdNT2/NLuc* reporter locus (flanked by both up- and downstream *LdNT2* IGRs) was amplified from transgenic parasites already harboring the plasmid depicted in A and subcloned into pBB for further manipulation. To ensure endogenous-like 5' processing, the preferred *LdNT2* splice acceptor site (SA, black circle) was maintained and integration was directed by the remaining upstream portion of 5'-IGR (preSA). C) Approach for deletional mutagenesis of the *LdNT1.1* 3'-UTR. The *LdNT1.1* 3' IGR was PCR amplified from genomic DNA as two separate segments, separated by the intended deletion site. Notably, constructs were transfected into a recipient cell line already expressing a firefly luciferase-blasticidin resistance fusion transgene (i.e. *Fluc-BSD)* from the endogenous *LdNT1.1* locus [Soysa, 2014], such that integration was directed by the *Fluc* reporter gene.
